# Extinction probabilities, times to extinction, basic reproduction number and growth rates for tsetse (*Glossina* spp) populations as a function of temperature

**DOI:** 10.1101/767350

**Authors:** Elisha B. Are, John W. Hargrove

**Author notes:** Private Bag X1 Matieland Stellenbosch, 7602.

## Abstract

Increases in temperature over recent decades have led to a significant reduction in the populations of tsetse flies (*Glossina* spp) in parts of the Zambezi Valley of Zimbabwe. If this is true for other parts of Africa, populations of tsetse may actually be going extinct in some parts of the continent. Extinction probabilities for tsetse populations have not so far been estimated as a function of temperature. We develop a time-homogeneous branching process model for situations where tsetse flies live at different levels of fixed temperatures. We derive a probability distribution *p*_*k*_(*T*) for the number of female offspring an adult female tsetse is expected to produce in her lifetime, as a function of the fixed temperature at which she is living. We show that *p*_*k*_(*T*) can be expressed as a geometric series: its generating function is therefore a fractional linear type. We obtain expressions for the extinction probability, expected number of female offspring per female tsetse, and time to extinction. No tsetse population can escape extinction if subjected, for extended periods, to temperatures outside the range 16 °C - 32°C. Extinction probability increases more rapidly as temperatures approach and exceed the upper and lower limits. If the number of females is large enough, the population can still survive even at high temperatures (28°C - 31°C). Small decreases or increases in constant temperature in the neighbourhoods of 16°C and 31°C, respectively, can drive tsetse populations to extinction. Further study is needed to estimate extinction probabilities for tsetse populations in field situations where temperatures vary continuously.

**Author summary:** Tsetse flies (*Glossina* spp) are the vectors of the African sleeping sickness. We derived an expression for the extinction probability, and mean time to extinction, of closed populations of the flies experiencing different levels of fixed temperatures. Temperatures play a key role in tsetse population dynamics: no tsetse populations can escape extinction at constant temperatures < 16°C > 32°C. The effect of temperature is more severe if tsetse populations are already depleted. Increasingly high temperatures due to climate change may alter the distribution of tsetse populations in Africa. The continent may witness local extinctions of tsetse populations in some places, and appearances in places hitherto too cold for them.

## Introduction

A bite from a tsetse fly (*Glossina spp*.) infected with a parasite of the genus *Trypanosoma* may cause Human African Trypanosomiasis (HAT), commonly called sleeping sickness in humans, or Animal African Trypanosomiasis (AAT), commonly called nagana in livestock. These tropical diseases have ravaged the African continent for centuries. They pose serious public health and socio-economic problems, especially to rural farmers, who rely on their livestock for daily subsistence, draught power and general economic gain. Sleeping sickness is difficult to diagnose and the treatments are often difficult to administer [1]. Vector control plays an important role in the fight against trypanosomiasis [2], and understanding the vector population dynamics is thus crucial.

As with all insects, the body temperature of tsetse is largely determined by ambient temperature and all of the flies’ physiological processes are determined by the temperatures that the flies experience. The flies use various behavioural devices to mitigate the effects of extreme ambient temperatures, such that the temperatures they actually experience are less extreme than indicated by temperatures measured in, for instance, a Stevenson screen [3, 4]. Nonetheless, excessively high, or low, temperatures may be lethal for them [5, 6]. This is a serious concern for tsetse because, unlike other insects, the genus *Glossina* is characterised by a very low birth rate. Consequently, small increases in mortality rates can result in negative growth rates that, if sustained, may drive tsetse populations to extinction.

As an example of this effect, a study published in 2018 concluded that, over the previous 40 years, there had been a significant increase in temperatures in the Zambezi Valley of Zimbabwe [7]. Specifically, peak temperatures at Rekomitjie Research Station in Mashonaland West Province, Zimbabwe, increased by c. 0.9°C from 1975 to 2017. The hottest time of the year (November) experienced a higher increase in temperature of c. 2°C during the same period. These increases in temperature may potentially explain the reduction in tsetse populations in some parts of Zimbabwe [8]. If these findings are true also for other parts of Africa, then it is important to estimate the impact of the high temperatures on the probability of, at least, local extinction of tsetse in some parts of the continent.

Two published works have estimated extinction probabilities and time to extinction for tsetse populations by assuming fixed environmental states throughout the life history of the flies. The first [9], derived a branching processes model for tsetse populations with the assumption that male and female offspring are produced with equal probability. This assumption is not generally true [10], although the results obtained in [9] are consistent with the body of the literature on tsetse biology. The second publication [11] provides a sound mathematical foundation for the results obtained in [9], and assumes a situation where male to female sex ratio can vary anywhere in the open interval (0, 1) and shows how extinction probability depends on the male-female sex ratio in tsetse populations.

In both papers, in order to gauge the importance of various determinants of tsetse population growth, extinction probabilities were calculated for numerous arbitrary combinations of mortality and fertility rates. There was no explicit modelling of the effect of temperature on vital rates, nor thus on its effect on extinction probabilities. In this paper, we develop a version of the stochastic branching process model presented in [9] and [11] in which all of the biological process in the tsetse lifecycle are explicitly dependent on temperature.

Various workers have obtained mathematical expressions for the temperature-dependence of such functions as daily survival probabilities for young, and mature, adult females, and pupae, and the inter-larval period and pupal duration [6, 8, 15–18]. Using these functions allows us to estimate extinction probabilities, time to extinction and mean of tsetse population size at different levels of fixed temperatures.

It was also assumed in the earlier papers that mortality in adult flies was independent of the age of the fly. Evidence suggests that mortality rates are actually markedly higher in recently emerged adult flies than in mature flies [6, 11–14], and that this difference is particularly severe at extremes of high temperatures [5, 6]. To capture this difference in mortality rates, we assume different survival probabilities for young adult flies for the first *ν* days after emergence, compared with the survival probability of all older flies.

## Materials and methods

We follow the general approach described earlier [9, 11] but the extinction probability is now obtained in a simpler form. We use the solution to obtain numerical results for extinction probabilities, time to extinction and growth rates for tsetse populations living at various fixed temperatures. We achieve this by using existing functions in the literature relating tsetse fly lifecycle parameters to temperature. We modify published versions of the model [9, 11] by separating the adult life stage of tsetse fly into immature and mature classes, allowing us to assess differential impacts of temperature in the two stages.

### Tsetse life history

The following provides a brief description of the life cycle; fuller accounts are provided in [19]. Unlike most other insects, tsetse have a very low birth rate: they do not deposit eggs, instead producing a single larva every 7 − 12 days [17, 20]. The larva buries itself in the ground and immediately pupates, staying underground as a pupa for between 30 − 50 days [21], emerging thereafter as an adult with the linear dimensions of a mature adult, but with poorly developed flight musculature, and low levels of fat. Both sexes of tsetse feed only on blood, and the first 2-3 blood-meals are used to build flight muscle and fat reserves, before the mature female can embark on the production of larvae of her own. All of these processes are temperature dependent. The pupal phase increases from 20 days at 32°C to 100 days at 16°C (28 days at 25°C) [22], though few adults emerge at the extremes of temperature. Blood-meals are taken every 2 – 5 days, again depending on temperature. The time between adult female emergence and first ovulation is only weakly dependent on temperature and is assumed here to take a constant value of 8 days in the field at Rekomitjie [19]. Thereafter, the period between the production of successive pupae increases from about 7 days at 32°C to 12 days at 20°C (9 days at 2° C) [20].

### Model

The model development and assumptions are similar to those in [11], differing only in the way that mortality and fertility rates are used in the models. In the earlier works, these rates were simply set at constant values. In the present study we use the fact that the rates are almost all known to be functions of the temperature experienced by the flies. Accordingly, instead of setting these rates at arbitrary values, we instead allow the environmental temperature to take various values, which then dictate the values of the rates of mortality and fertility to be used in the model. In particular, the following rates are all temperature dependent; pupal duration and inter-larval period, and the daily survival probabilities for female pupae, adult females that are immature (defined as not having yet ovulated for the first time), and mature adult females.

### Model assumptions

In what follows all parameters with subscript *T* are temperature dependent.

1. An immature adult female tsetse fly [6] survives the *ν* days until it ovulates for the first time with probability 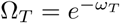 mortality rate. per day, where *ω*_*T*_, is the daily
2. A mature female tsetse survives with probability 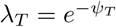; per day, where *ψ*_*T*_, is the daily mortality rate for mature females.
3. Once a female has ovulated for the first time, she deposits a single larva every *τ*_*T*_ days.
4. A deposited larva is female with probability *β*.
5. The larva burrows rapidly into the soft substrate where it has been deposited and pupates [8]. The pupa survives with probability 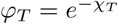 per day; where *χ*_*T*_ is the daily mortality rate for female pupae.
6. At the end of the pupal period of *σ*_*T*_ days, an immature adult fly emerges from the puparium.
7. The immature female fly is inseminated by a fertile male tsetse after *ν* days, with probability *ϵ*.

The probability that a female tsetse produces *k* surviving female offspring is obtained as:

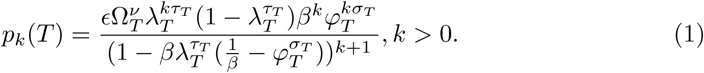

The proof of equation (1) can be easily adapted from the proof of equation (7) in the supplementary material of [11]. The difference here is simply that the probability is a function of temperature. We assume that *T* is time invariant. Our interest is to estimate extinction probabilities for tsetse at fixed temperatures.

Equation (1) can be used to derive a function for the extinction using the procedures described in earlier publications [9, 11]. A simpler derivation, using the work of Harris [22], is given here. Suppose *p*_*k*_(*T*) follows a geometric series for all *T*, then:

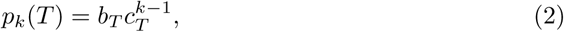

where *b*_*T*_, *c*_*T*_ > 0.

For equation (1) we then have:

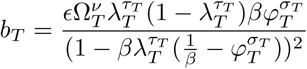

and

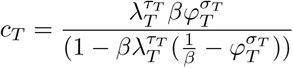

Inserting *b*_*T*_ and *c*_*T*_ into equation (2), yields

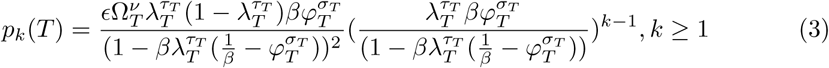

It follows (from [22], page 9) that the generating function *f*_*T*_ (*s*), for *p*_*k*_(*T*) is a fractional linear function, and can be expressed as:

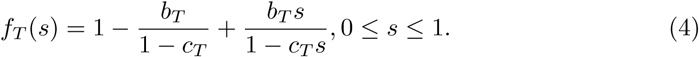

### Mean of female tsetse population at generation n

Substituting for *b*_*T*_ and *c*_*T*_ in equation (4) and taking the first derivative w.r.t s, at *s* = 1.

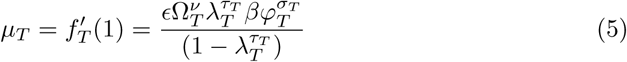

Equation (5) is the reproduction number for a female tsetse population. For a population of tsetse living at temperature T°C, the expected number of female tsetse in the population at generation *n* is denoted by

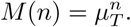

#### Remark 1

When *µ*_*T*_ > 1, the branching process is said to be supercritical with extinction probability *q*_*T*_ < 1. If *µ*_*T*_ < 1, the branching process is subcritical, which implies, in practice, that each female tsetse produces less than one surviving female offspring on average. Extinction is then certain: i.e., the probability *q*_*T*_ = 1. The process is called critical if *µ*_*T*_ = 1, and extinction probability is again certain, *q*_*T*_ = 1 (see [23] page 36). In other words, for any tsetse population to avoid inevitable extinction, each female fly must produce more than one surviving female offspring in her lifetime.

### Extinction probability *q*_*T*_

The extinction probability is obtained by solving for the fixed points of equation (4), that is, we find s such that *f*_*T*_ (*s*) = *s*. We therefore need to solve:

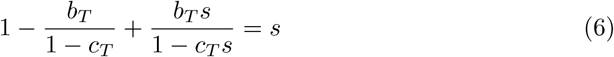

Substituting for *b*_*T*_ and *c*_*T*_ in equation (6) and solving for *s*, the extinction probability *s* = *q*_*T*_ is the smaller nonnegative root of equation (6).

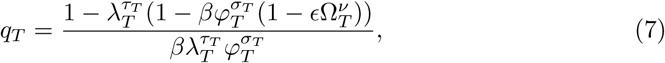

where 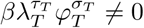.

#### Remark 2

Suppose 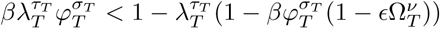, then 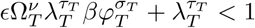, which implies that *µ*_*T*_ < 1 (5)). Therefore, whenever the denominator of equation (7) is less than the numerator, extinction probability *q*_*T*_ = 1. Hence, for all biologically meaningful parameter ranges, *q*_*T*_ is always in [0, 1]. See **Remark 1**. Furthermore, when the initial population consists of a single female fly, the extinction probability is given by equation (7). If the initial population is made up of *N* female flies, the extinction probability is given by (*q*_*T*_)^*N*^ [11].

### Expected time to extinction for a population of tsetse

An expression for the expected time for population of the female tsetse to become extinct is presented in [9], as:

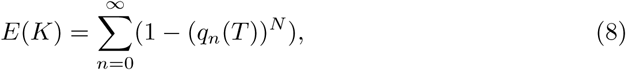

where

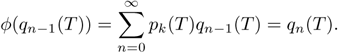

*K* is the time to extinction and *N* is the number of female tsetse flies in the initial population. To estimate the mean time to extinction of a population consisting of *N* flies, it suffices to calculate *q*_*n*_(*T*) iteratively, and raise each value to the power *N*, to obtain *E*(*k*) as given in equation (8). Note that, since females produce both male and female offspring, extinction of the female population obviously guarantees the extinction of the whole population.

### Tsetse mortality rates as a function of temperature

The relationship between temperature and the instantaneous daily mortality rate of pupae is modelled as a the sum of two exponentials (Fig1) [15].

**Fig 1.**
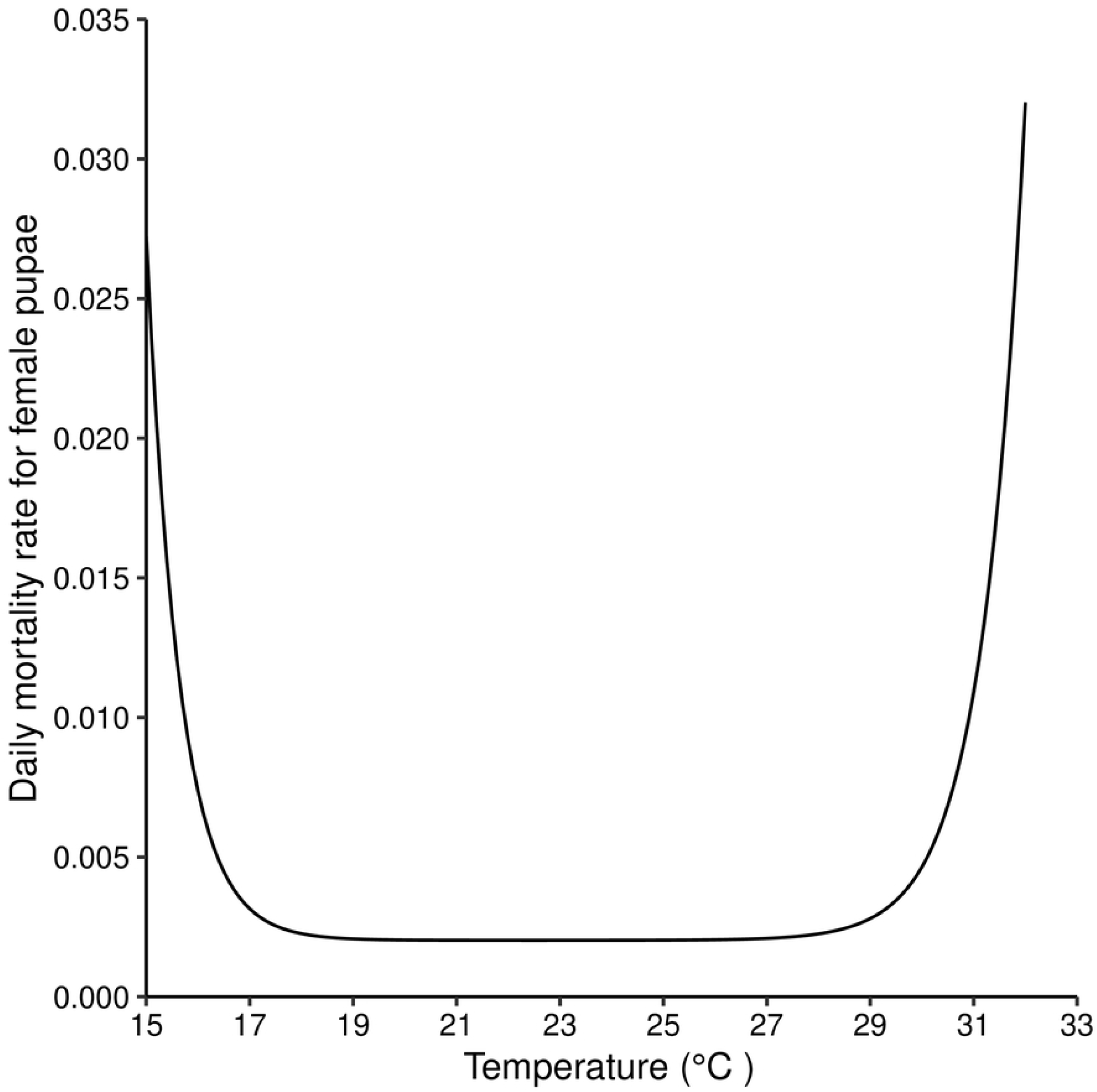
Daily mortality rates for female *G. m. morsitans* pupae for temperatures between (15°C −33°C). Equation (9) plotted for different values of temperature.

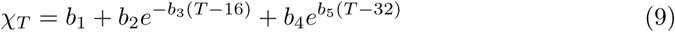

Daily mortality rates of young and mature adult flies increase exponentially with temperature (Fig2). The defining equations, (10) and (11), take the same form, differing only in the parameter values (Table 1), due to the higher mortality of young flies, particularly at high temperatures.

**Fig 2.**
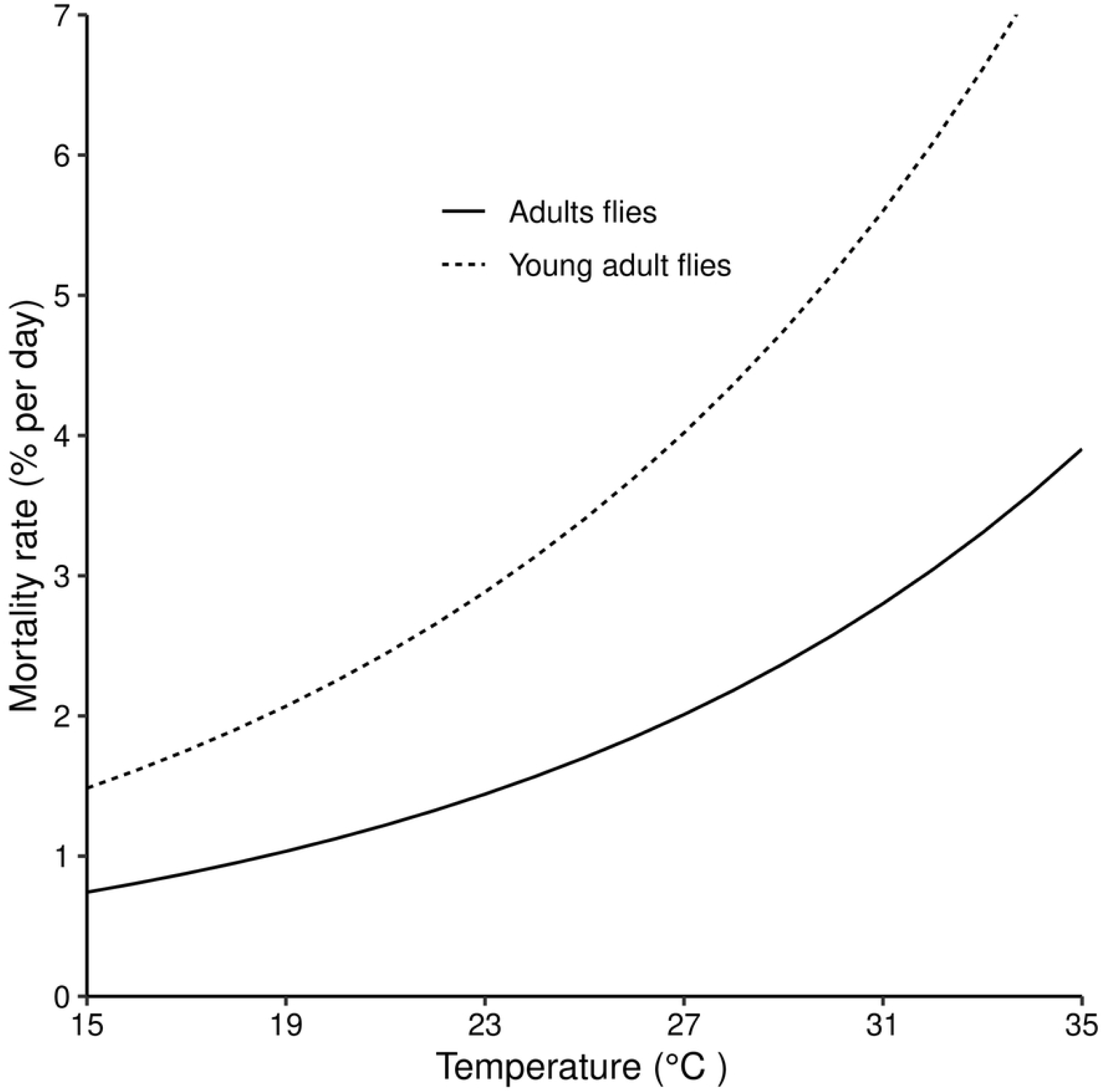
Daily mortality rates for young adult and mature adult female tsetse for temperatures ranging from 15°C - 35°C. Equations (10) and (11) plotted for different temperatures.

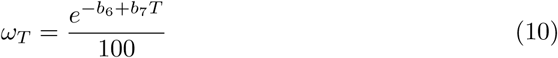

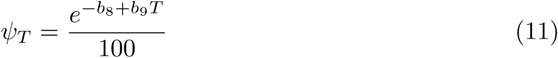

**Table 1.** Summary of model parameter values and sources: Parameter values are generally sourced from the literature.

| Parameter | Description | Value & Source |
| --- | --- | --- |
| $b_1$ | Daily female pupal mortality ($\chi_T$) | [15]<br>0.00202 |
| $b_2$ | Equation (9) | 0.00534 |
| $b_3$ | | 1.55200 |
| $b_4$ | | 0.03000 |
| $b_5$ | | 1.27100 |
| $b_6$ | Daily mortality rate for young flies ($\omega_T$) | [18]<br>-0.85000 |
| $b_7$ | Equation (10) | 0.08300 |
| $b_8$ | Daily mortality rate for mature flies ($\psi_T$) | [18]<br>-0.15430 |
| $b_9$ | Equation (11) | 0.08300 |
| $c_1$ | Pupal duration ($\sigma_T$) (days) | [17]<br>17.94000 |
| $c_2$ | Equation (12) | 82.30000 |
| $c_3$ | | -0.25300 |
| $d_1$ | Inter-larval period ($\tau_T$) (days) | [18]<br>0.10460 |
| $d_2$ | Equation (13) | 0.00520 |
| $\beta$ | Probability deposited pupa is female $\beta$ | 0.5 [11] |
| $\nu$ | Time (days) from adult emergence to first ovulation | 8.0 [11] |
| $\epsilon$ | Probabilty of insemination by a fertile male | 1.0 [11] |

### Development rates as a function of temperature

Pupal duration is modelled as increasing exponentially with decreasing temperature ([15], [24–26]).

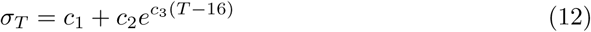

The relationship between larviposition rate and temperature was modelled using the results from a mark and release experiment conducted at Rekomitjie on *G. m. morsitans* and *G. pallidipes* [17].

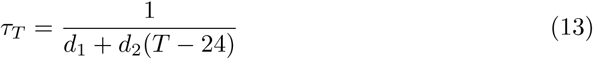

In equations (9) – (13), *b*_1_, *b*_2_, *b*_3_, *b*_4_, *b*_5_, *b*_6_, *b*_7_, *b*_8_, *b*_9_, *c*_1_, *c*_2_, *c*_3_, *d*_1_ and *d*_2_ are all constants.

Numerical simulations were carried out using R studio (version 1.1.463) [27], by incorporating equations (9) – (13) into equation (7) and taking parameter values from the literature as shown in Table 1.

## Results

### Extinction probability as a function of fixed temperatures

Extinction probabilities were calculated as a function of fixed temperatures ranging from 15 – 35°C, for different values of the number of females in the initial population. As expected, sustained extreme temperatures can drive tsetse populations to extinction. When the initial population consists of a single female fly, extinction probability did not drop below 0.45 even with optimal temperatures (Fig3). However, when the starting population consists of even 10 female flies, extinction probability drops rapidly to zero as temperatures increase slightly above 17°C.

**Fig 3.**
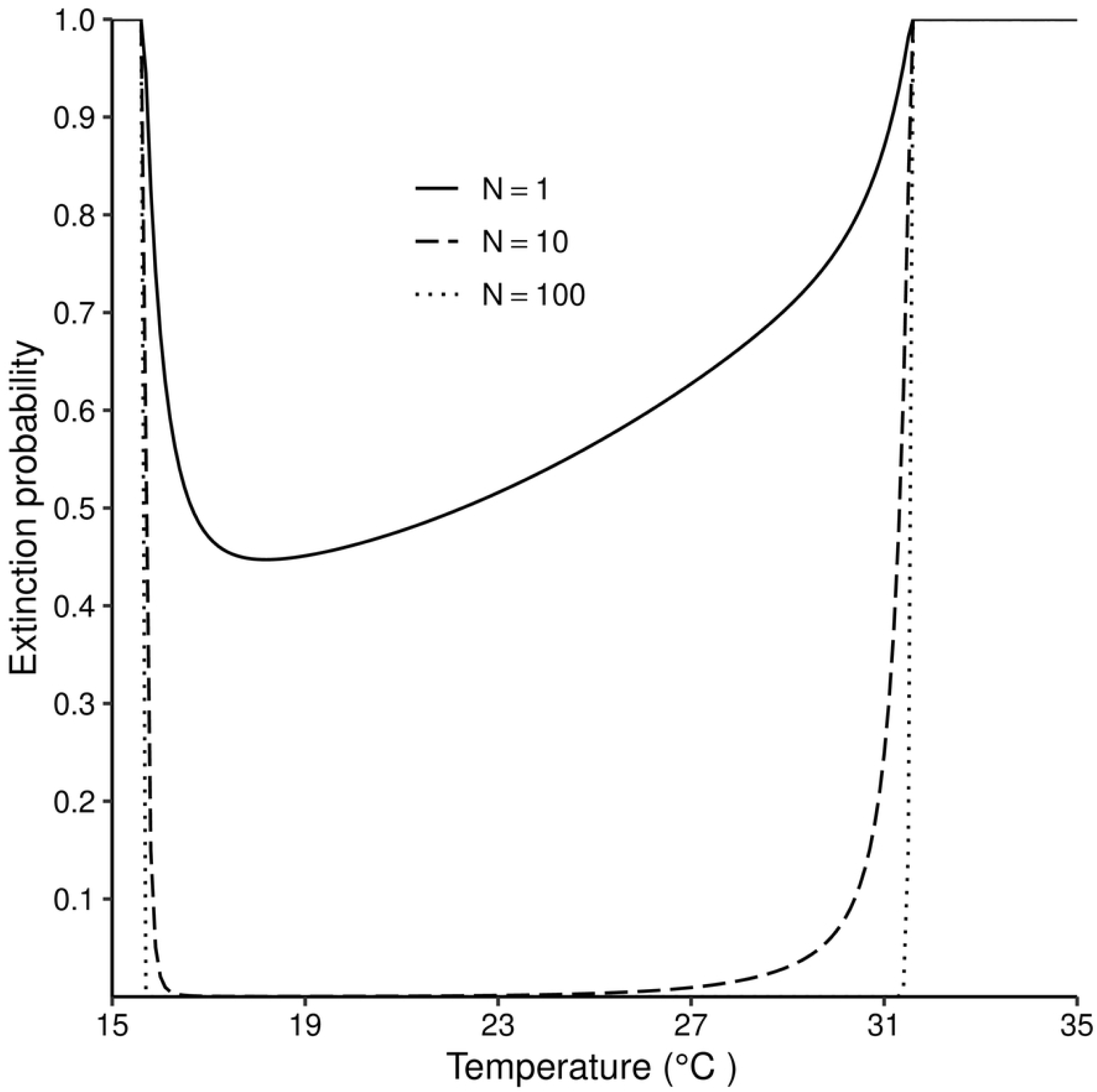
Probability of extinction as a function of temperature varying between 15°C and 35°C. Equation(7) solved for different values of temperature and for different numbers of inseminated adult females in the initial population.

With sufficient female tsetse, say 100, in the initial population, extinction probabilities went rapidly to zero when temperatures exceeded 16°C, and remained at 0 even at temperatures slightly above 31°C. No tsetse population, regardless of the number of females in the starting population, can escape extinction outside the range of approximately 16°C - 32°C (Fig3).

### Expected number of females per female tsetse for different levels of fixed temperatures

The reproduction number for a population, i.e., the number of surviving daughters an adult fly is expected to produce in her lifetime, is presented in Figure 4 for different levels of fixed temperatures. The reproduction number peaks at 2.8, for temperatures in the neighbourhood of 19°C. It gradually declines as temperatures increase above 21°C, dropping below 1.0 when temperatures exceed about 31°C. However, as temperature decreases below 18°C there is a sharp drop in the reproduction number, which goes below 1.0 at a temperature just below 16°C.

**Fig 4.**
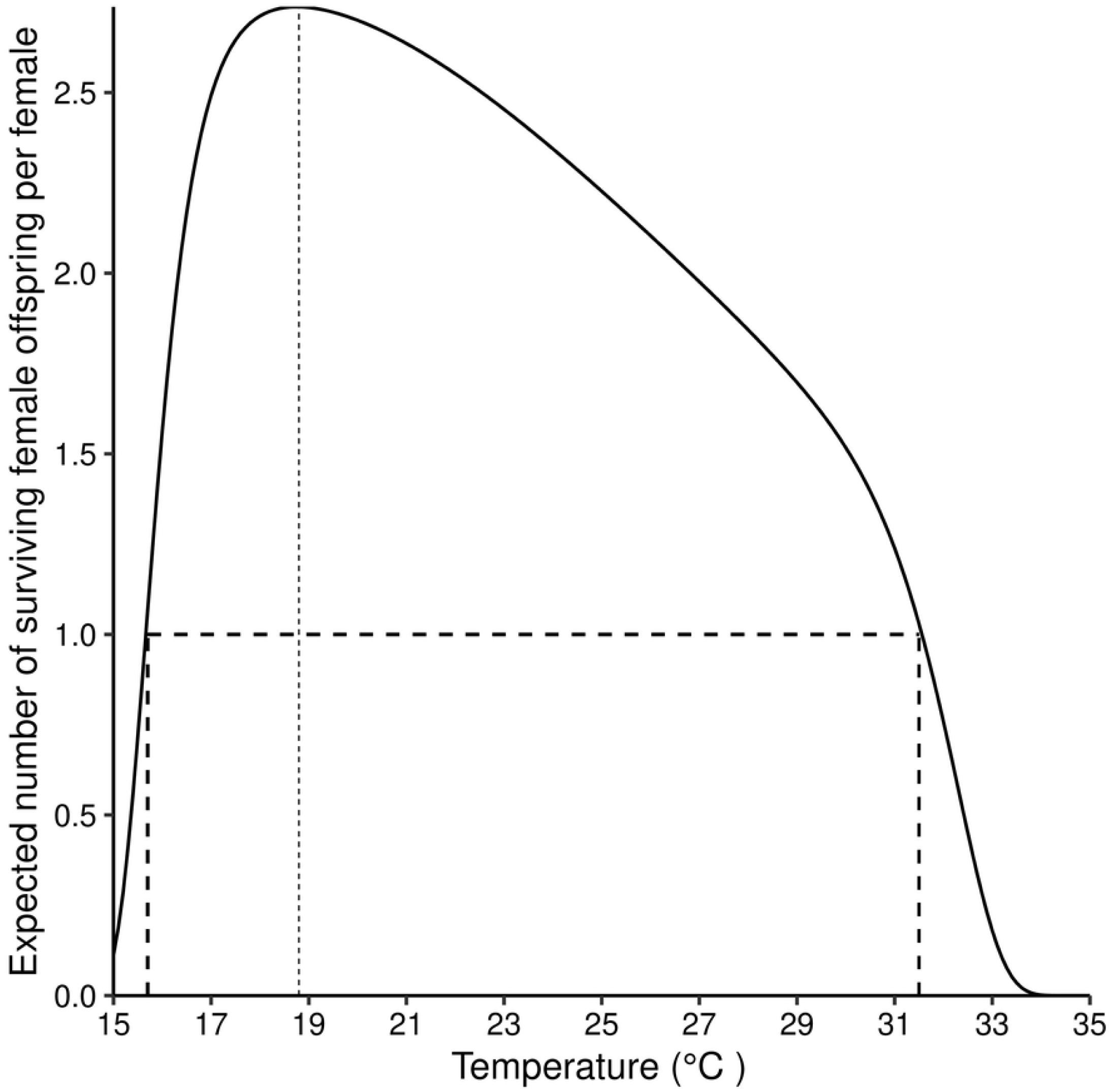
Expected number of surviving female offspring per adult female tsetse for different temperatures (15°C-35°C). Equation(5) solved for different values of temperature.

### Time for the population of female tsetse flies to go extinct at different fixed temperatures

The expected number of generations to extinction varies with temperature and with the size of the starting population. When the simulation for the expected number of generations is done for 20 generations, it takes 12 generations for a population starting with 2 female flies to go extinct under optimal temperatures of 19 - 21°C. As temperature increases above 21°C, the number of generations to extinction reduces gradually, but the decrease becomes much more rapid as temperatures approach 31°C When the number of flies in the initial population increases to 10, extinction did not occur in the first 20 generations for temperatures between 16 and 31°C (Fig5).

**Fig 5.**
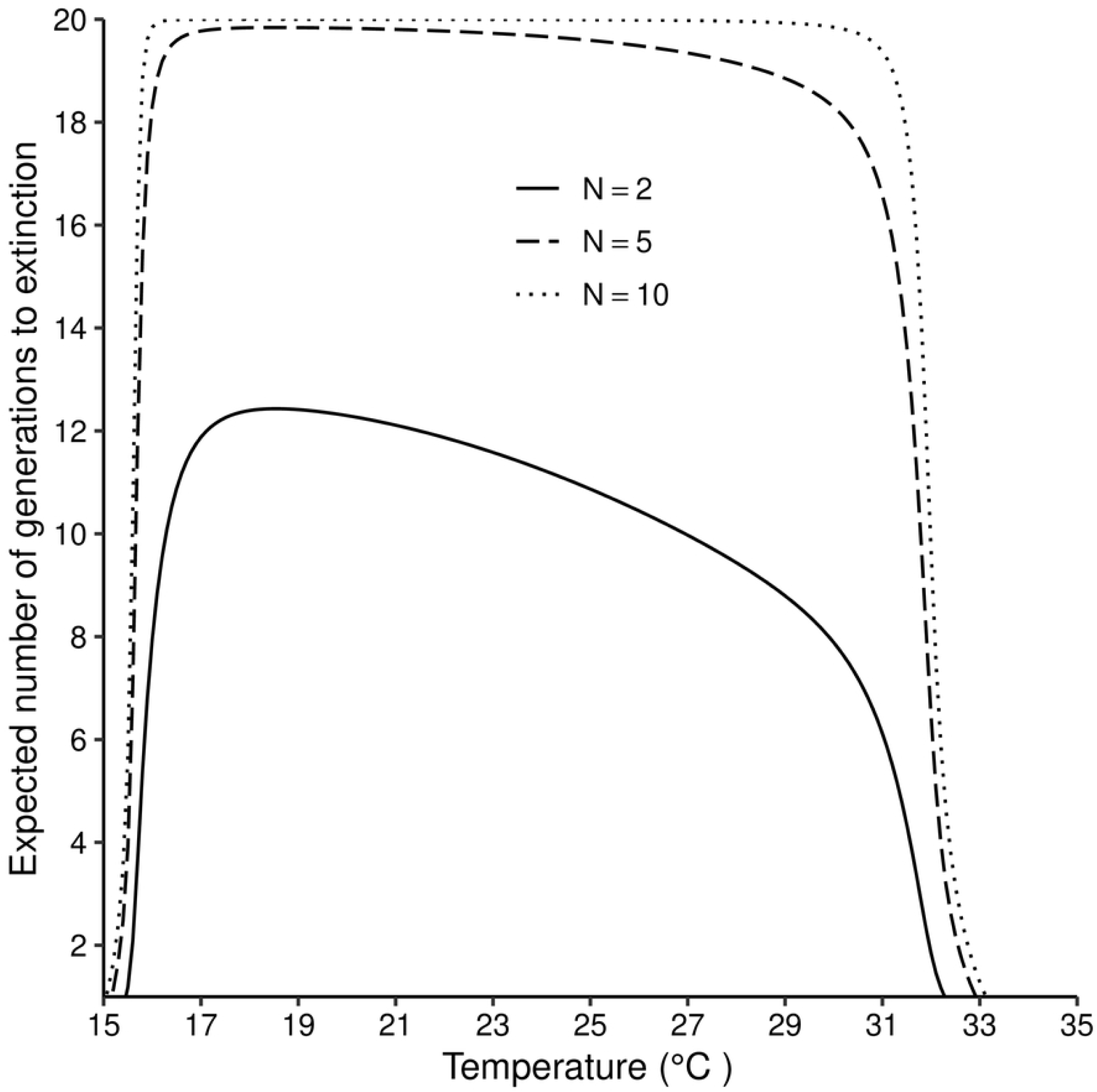
Expected number of generation to extinction for different number of female in the initial population at different temperatures (15°C-35°C). Equation (8) is solved iteratively up to *n* = 20.

### Tsetse population growth rates at different fixed temperatures

Tsetse populations, which are small enough that we may ignore density dependent effects, grow exponentially for temperature in the approximate range 16 - 31°C (Fig6). Notice that, in this figure, growth is plotted as a function of the number of generations completed. In order to gauge the growth rate as a function of time it is necessary to adjust for the fact that the generation time increases with decreasing temperature.

**Fig 6.**
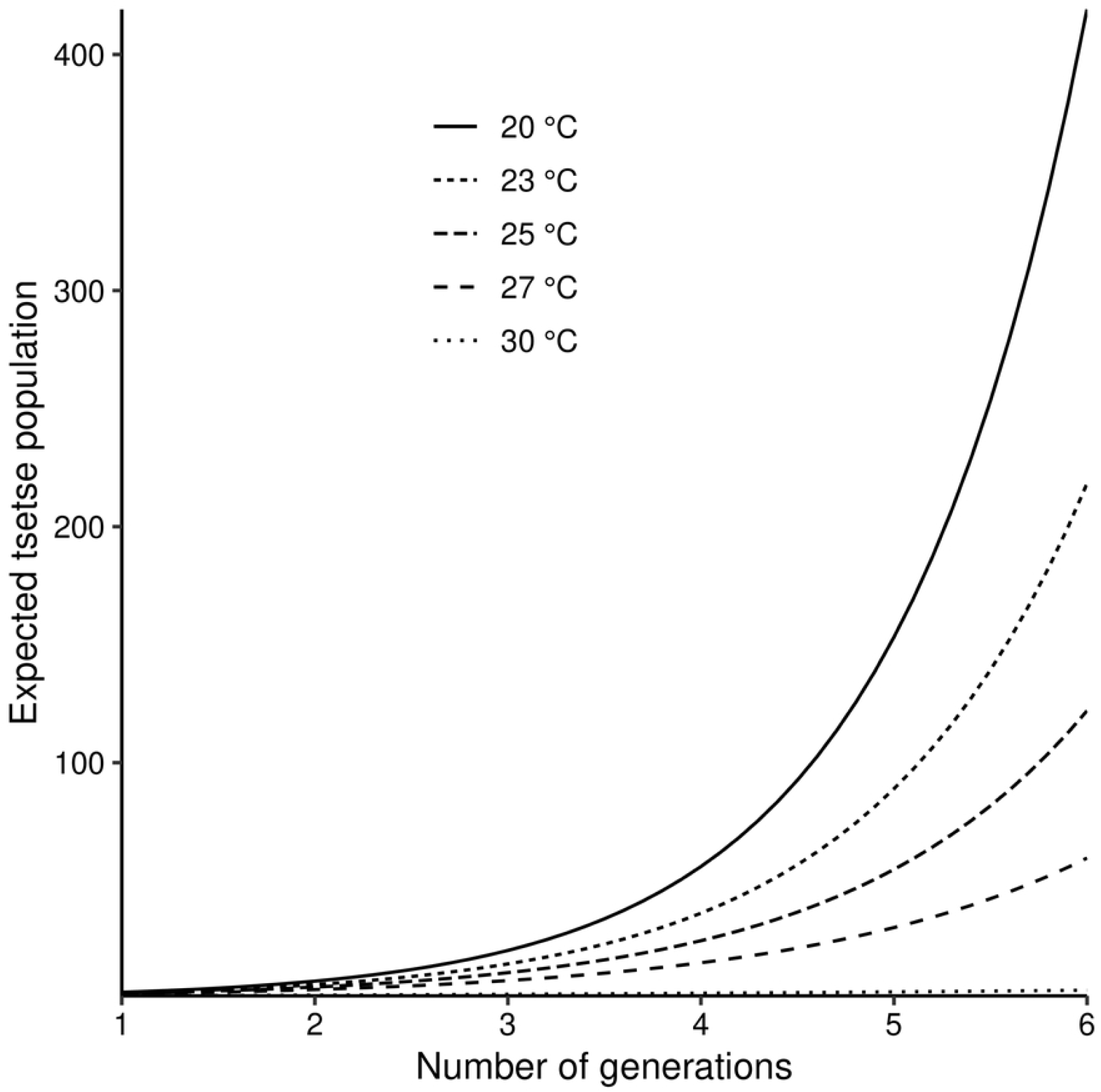
Expected growth in the numbers of adult females in a tsetse population at different temperatures (15°C-35°C). The projections are approximately valid for the early stages of growth, before density dependent processes have noticeable effects.

After controlling for generation time, the growth rate of the population as a function of calendar time is seen to take a maximum value of about 2.0% per day at 25°C, falling away increasingly rapidly towards zero as temperatures approach the upper and lower limits of 32°C and 16°C, respectively (Fig7).

**Fig 7.**
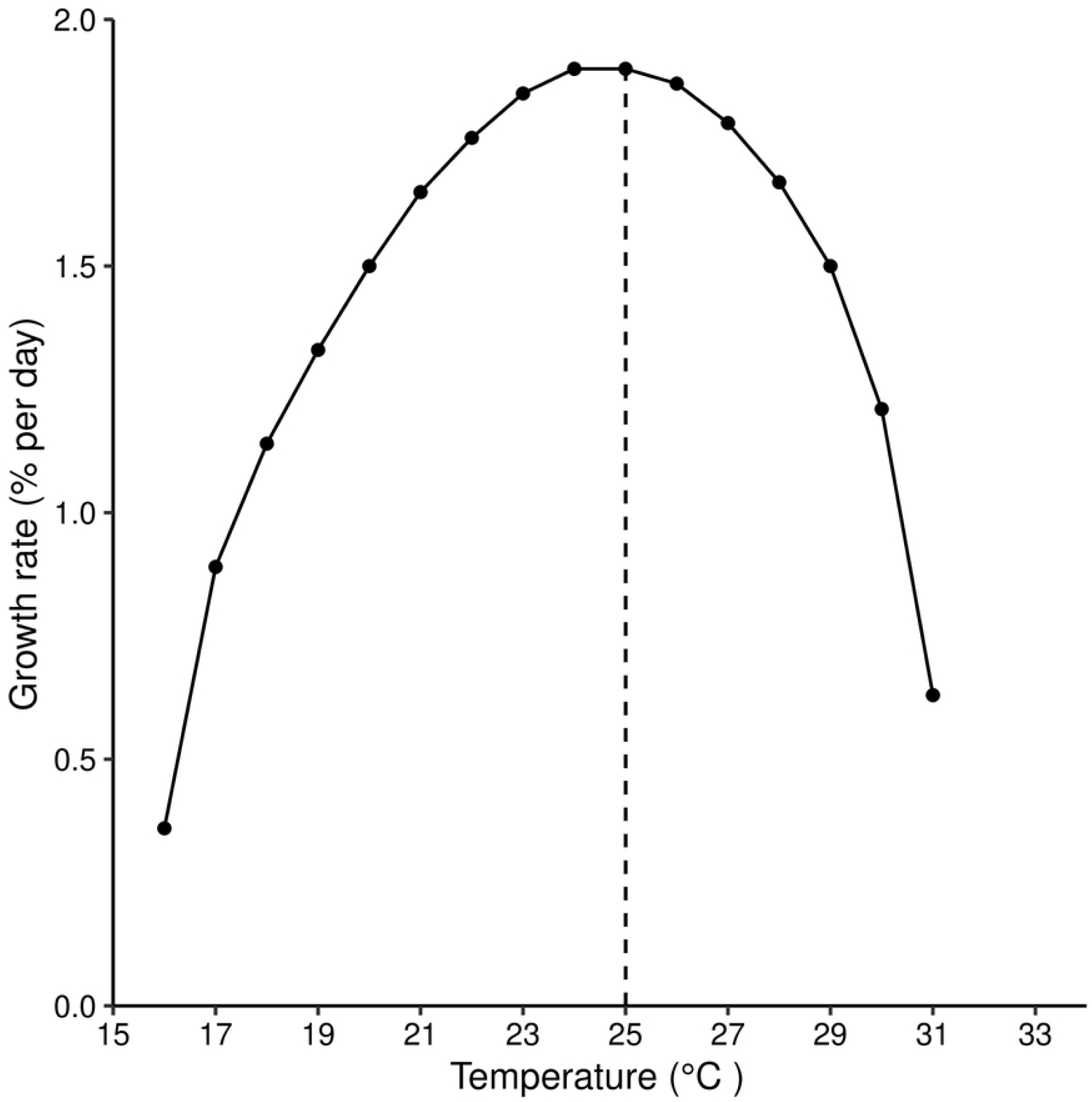
Population growth rate per day (%) at different temperatures (15°C-35°C). After controlling for generation time at different temperature levels.

## Discussion

The aim of this study was to develop a branching process model for tsetse populations experiencing fixed temperatures of different levels, similar to laboratory situations where tsetse flies are kept under regulated temperatures. We estimate extinction probabilities, times to extinction, expected numbers of female offspring per individual female, and growth rates for each scenario. This enables us to determine optimal temperatures for reproduction, and the lower and upper bound temperatures for the survival of tsetse populations.

Our results confirm findings of earlier studies which suggest that temperature is a key driver of tsetse population dynamics [8, 28, 29]. We show that constant temperatures outside the approximate bounds of 16 - 31°C are fatal for any tsetse population. These results are consistent with observations on tsetse populations in the Zambezi valley of Zimbabwe, where extreme temperature events have led recently to significant reductions in tsetse populations [8].

As temperatures increase, mortality rates increase for adult female tsetse, and for pupae of both sexes [26, 30], but larval production rates also increase and pupal durations decline [18]. In this trade-off, extinction probability initially declines as temperature increase above 15°C (Fig2). In fact even for quite small pioneer populations the extinction probability falls rapidly to zero for temperatures between 17 and 27°C. Thereafter, however, increases in mortality rates outweigh the increases in birth rates and the extinction probability increases ever more rapidly as temperatures approach 32°C.

When temperatures approach the lower limit of the 16°C– 32°C bracket, adult temperature-dependent mortalities decline to low levels, but female tsetse reproduction rates fall drastically as pupal durations increase. In the extreme, when temperatures drop below 16°C, pupal durations are so long that fat reserves are exhausted before the pupa can emerge [18]. The combination of these factors, if sustained, ensures extinction of tsetse population at very low temperatures (Fig2).

Theoretically, for any population to be able to escape extinction, each female adult must produce strictly more than one female offspring, which must themselves survive to reproduce. In epidemiological terms, we then have that the reproduction number, *R*_*o*_ > 1. Figure4 shows that for temperatures below 16°C, female tsetse will not be able to produce enough female flies to sustain the population. If the cold temperature conditions are prolonged, *Ro* will drop below 1, resulting ultimately in extinction. Thus, temperatures approach either hot or cold limits, the number of generations that a population can survive goes rapidly to zero even for large pioneer populations (Fig4).

The reproduction number reached its highest value of 2.8 at 19°C (Fig4). This may, initially, suggest that the growth rate is highest at that temperature. However, when we calculated the actual growth rate after controlling for the length of generation for different temperature values, we found that the population attains its maximum growth rate at 25°C. This result agrees with published values in the literature [18], where a different method was used to obtain the same results. It also draws attention to the fact that the reproduction number should be used with caution when comparing two populations with differing lengths of generation.

Our findings are in good agreement with experimental, field and modelling studies of the impact of temperature on different tsetse species [8, 28, 31–33], For instance, in [33] an experiment was conducted on three different strains of *Glossina palpalis gambiensis*, in a bid to determine critical temperature limits for tsetse survival and their resilience to extreme temperatures. For the three strains, a temperature of about 32°C was reported as the upper limit of survival. Our results showed, similarly, that, if the number of female flies in the population are high enough, tsetse population may escape extinction at temperatures slightly above 31°C but will go extinct at higher temperatures. The experiment also reached the conclusion that temperatures of about 24°C are optimal for rearing this species, in good agreement with our modelling results.

### Limitations of the study

Extinction probabilities for tsetse populations have not previously been estimated as a function of temperature. Our modelling framework took into consideration the fact that, as poikilotherms, tsetse mortalities, development rates, larval deposition rates, are all temperature dependent. However, the present study did not consider field situations where temperatures vary continuously with time. A modelling framework which will consider this more realistic situation is under construction.

In this study, we considered closed tsetse populations, where there was no in-or-out migration. Estimation of extinction probabilities for populations that are open to migration are markedly more complex and were beyond the scope of the present study. We do note, however, that a preliminary study found that if tsetse populations are in patches, which can compensate for each other, then extinction probabilities are lower than in closed population models [34]. Further work in this area is called for.

Finally, we caution that the modelling here is restricted to situations where population numbers lie below the level at which density dependent effects play a significant role. When numbers are larger, however, suppose pupal and/or adult mortality increases with density. Then the temperature-dependent mortalities quoted here provide a lower limit of the true mortality at that time, for any given temperature. Similarly, if density dependence results in a decrease in the birth rate, then the birth rates quoted here provide an upper bound to the true birth rate. For such a population, it then follows that the growth rate at any temperature will be lower than calculated here. Moreover, as temperatures increase, or decrease, towards the upper or lower bounds, respectively, for positive population growth, the population will decline rapidly towards levels where the density-dependent effects fall away. At that point the further dynamics of population growth would be subject to the vital rates used in this study.

